# Adaptation of white adipocytes to cooler temperatures: impacts on energy metabolism and protein acetylation

**DOI:** 10.64898/2026.04.14.718465

**Authors:** Hiroyuki Mori, Hadla Hariri, William Moe, Sophia Durham, Yuridia Guzman, Emma Paulsson, Rachel C. Simmermon, Parth B. Bhanderi, Sydney K. Peterson, Mia J Dickson, Charles R. Evans, Ormond A. MacDougald

**Affiliations:** Department of Molecular & Integrative Physiology, University of Michigan Medical School, Ann Arbor, Michigan, USA; Department of Internal Medicine, University of Michigan Medical School, Ann Arbor, Michigan, USA

**Keywords:** Adipocytes, Cool-temperature adaptation, Mitochondrial function, Protein acetylation, Metabolic remodeling, SHMT2, PCCA

## Abstract

Adipocytes throughout the body reside in distinct thermal environments. Visceral adipocytes within the body core are maintained near 37 °C, whereas those in bone marrow, subcutaneous, and dermal depots occupy cooler regions within the peripheral shell. While brown and beige adipocyte responses to cold stress are well characterized, much less is known about how white adipocytes adapt to moderately reduced temperatures below 37 °C. Our recent work revealed that cultured adipocytes exposed to 31 °C, a temperature representative of distal adipose regions, exhibit enhanced mitochondrial function, including increased substrate oxidation and ATP turnover, yet the mechanisms underlying this upregulation remain unclear. Here we show that adaptation to cool temperatures leads to a widespread decrease in protein acetylation in both undifferentiated and differentiated adipocytes, independent of nutrient status, and that this change is readily reversible upon rewarming. Subcellular fractionation and immunoblotting demonstrate that the hypoacetylation coincides with a compartment-specific enrichment of acetylated proteins within mitochondria, indicating selective remodeling of the mitochondrial acetylome. Transcriptomic and biochemical analyses reveal that these temperature-dependent changes occur without alterations in acetyltransferase or deacetylase expression, NAD⁺ concentration, or acetyl-CoA availability, suggesting regulation through alternative mechanisms affecting acetyl-CoA flux or enzyme activity. Integrative acetyl-proteomic and metabolomic profiling identifies mitochondrial enzymes, including serine hydroxymethyltransferase 2 (SHMT2) and propionyl-CoA carboxylase α (PCCA), whose acetylation correlates closely with changes in associated metabolite pools. Together, these findings establish physiologically relevant cooling as a cell-autonomous regulator of mitochondrial protein acetylation and metabolic adaptation in adipocytes.

## INTRODUCTION

Temperature is a fundamental yet often underappreciated regulator of cell biology. In homeothermic organisms, core body temperature is maintained near 37°C, but substantial thermal heterogeneity exists across tissues: peripheral extremities, subcutaneous depots, skin, and bone marrow frequently operate several degrees cooler than the body core, whereas highly active organs can be locally warmer than ambient or core temperature [1; 2]. Adipose depots exemplify this variability; subcutaneous, dermal, and marrow adipocytes typically experience cooler microenvironments than visceral adipocytes located in the abdominal cavity. This spatial temperature gradient raises the possibility that the same cell type may adopt distinct metabolic and signaling states depending on its thermal niche [1; 2].

Recent studies have shown that temperature changes *in vitro* can autonomously drive significant shifts in cell phenotype and function. For example, culturing adipocytes at cooler temperatures induces thermogenic genes such as UCP1 and promotes browning, independent of traditional adrenergic signaling pathways [3; 4]. Building on these observations, recent work has shown that white adipocytes cultured at 31°C, which closely matches distal bone marrow and subcutaneous tissue temperatures, undergo a marked remodeling of their metabolic program. Compared with 37°C, cool-adapted adipocytes use less glucose and instead rely more heavily on pyruvate, glutamine, and fatty acids as energy substrates, accompanied by enhanced lipid turnover and improved mitochondrial function, including increased mitochondrial content and higher respiratory capacity [5]. These data indicate that modest cooling, without overt cold stress, is sufficient to rewire substrate preference and oxidative metabolism in adipocytes. However, the molecular mechanisms that couple even a small decrease in temperature to such pronounced changes in mitochondrial activity remain poorly defined.

Acetylation of proteins on lysines is a post-translational modification that links cellular metabolism to enzyme function, organelle activity, and gene regulation [6]. In mitochondria, acetylation of metabolic enzymes can alter catalytic efficiency, substrate channeling, and pathway flux, while in the nucleus and cytosol, acetylation modulates transcriptional programs and signaling networks [7; 8]. Because acetylation depends on acetyl-CoA availability, redox state, and deacetylase activity, it is well positioned to act as a sensor–effector system that integrates environmental cues, including temperature, with metabolic output [7; 8]. Yet, how physiologically relevant “cool” temperatures influence the global acetylation landscape, and whether temperature-dependent acetylation of specific mitochondrial enzymes contributes to the metabolic remodeling of adipocytes, has not been systematically examined.

Previous work demonstrated that adipocytes cultured at 31°C exhibit elevated mitochondrial respiration, increased β-oxidation, and upregulated OXPHOS complex protein levels [5]. Here, we test the hypothesis that temperature-dependent changes in protein acetylation contribute to this upregulation. Using cultured adipocytes, we examine how shifting cells from 37°C to 31°C affects global and compartment-specific protein acetylation, with particular focus on mitochondrial proteins. We then interrogate classical regulators of acetylation, including lysine acetyltransferases, deacetylases, and acetyl-CoA metabolism, to determine whether these can account for the observed temperature-dependent changes. Finally, by integrating acetyl-lysine proteomics with targeted metabolomics, we identify candidate enzymes, such as serine hydroxymethyltransferase 2 (SHMT2) and propionyl-CoA carboxylase alpha chain (PCCA), whose acetylation states correlate with altered metabolite pools. Together, these approaches aim to define how modest cooling acts as a regulator of non-histone protein acetylation and to reveal temperature-sensitive regulatory points in mitochondrial metabolism that may underlie the adaptive phenotype of cool-adapted adipocytes.

## RESULTS

### Cool temperatures enhance mitochondrial respiration ex vivo

We have previously demonstrated that adipocytes cultured at 31°C exhibit elevated mitochondrial OXPHOS capacity by Seahorse XF analyses (Complex I, II, III-linked respiration) and upregulated protein levels of OXPHOS proteins [5]. To assess changes in mitochondrial respiration *ex vivo*, we measured bioenergetic parameters of posterior subcutaneous white adipose tissue (ps-WAT)-derived floated adipocytes isolated from mice housed at either 30°C or 22°C for 12 weeks. Isolated adipocytes exhibited a significant increase in baseline respiration, maximum respiration, and spare capacity upon Oroborus Oxygraph-2k analyses (**Figure 1A, 1C, 1D**). ATP-linked respiration was unchanged between temperatures (**Figure 1B**). Our data suggest that cool temperatures alter non-ATP-consuming oxygen utilization such as proton leak and substrate oxidation, enhance mitochondrial oxidative potential, and improve mitochondrial adaptability to respond to energy stresses. At the molecular level, little is known about the mechanisms underlying enhanced mitochondrial respiration and remodeling at cool temperatures.

**Figure 1.**
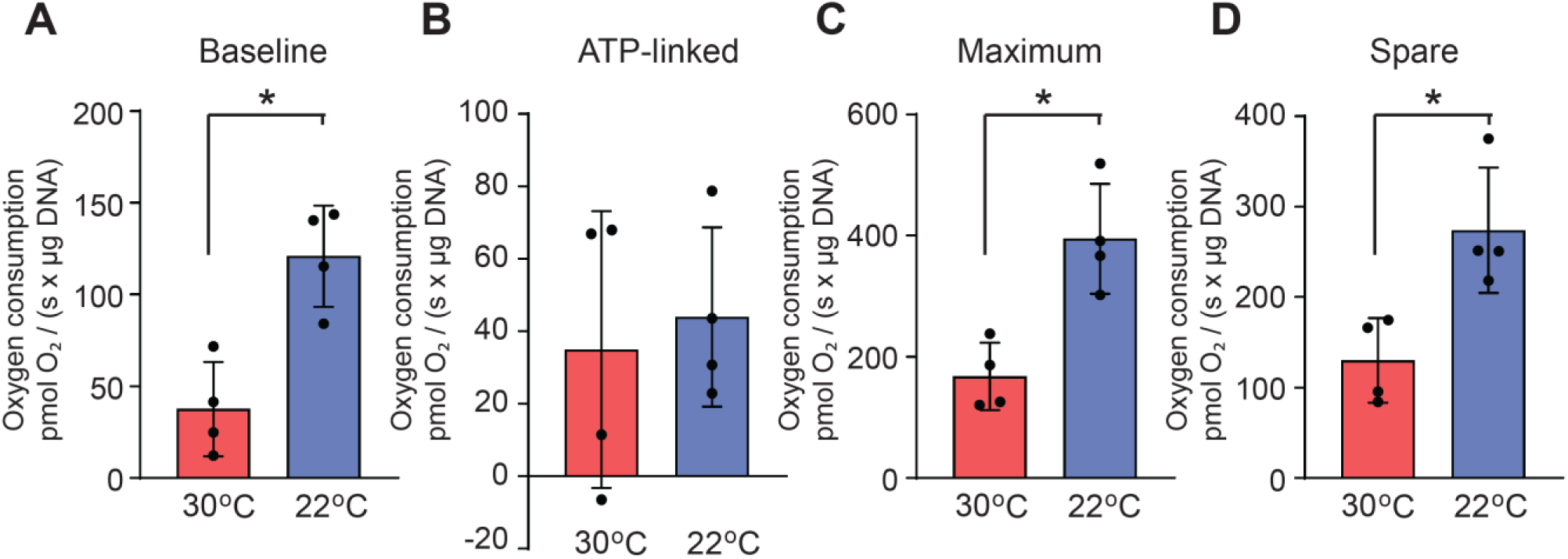
Housing mice at 22°C enhances mitochondrial respiration *ex vivo.* Oroboros Oxygraph 2k analyses of isolated adipocytes from psWAT depots of male mice at an environmental temperature of either 30°C or 22°C for 12 weeks. (**A**) Baseline respiration, (B) ATP-linked respiration, (**C**) maximum respiration, (**D**) spare capacity (*n=4*). Data are presented as mean ± SD. \**p* < 0.05.

### Cool adaptation reduces global protein acetylation in mesenchymal stem cells and differentiated adipocytes, independent of nutrient status and is reversible

To investigate effects of cooler temperatures on protein acetylation, adipocytes were cultured at 31°C for specified intervals (**Figure 2A**). Immunoblotting for acetylated lysine revealed a pronounced reduction in protein acetylation following exposure to 31°C, most notably after 8 days. Because acetyl-CoA is a key substrate for protein acetylation and its cellular abundance can be affected by presence of nutrients and growth factors [7; 9], cells were incubated in media containing either 2% FBS for 4 hours or 10% FBS for 16 hours prior to harvest. Importantly, adipocytes exposed to 31°C exhibited marked decreases in acetylated protein levels regardless of FBS amount, demonstrating that cool-induced hypoacetylation is independent of external serum concentrations (**Figure 2A**).

**Figure 2.**
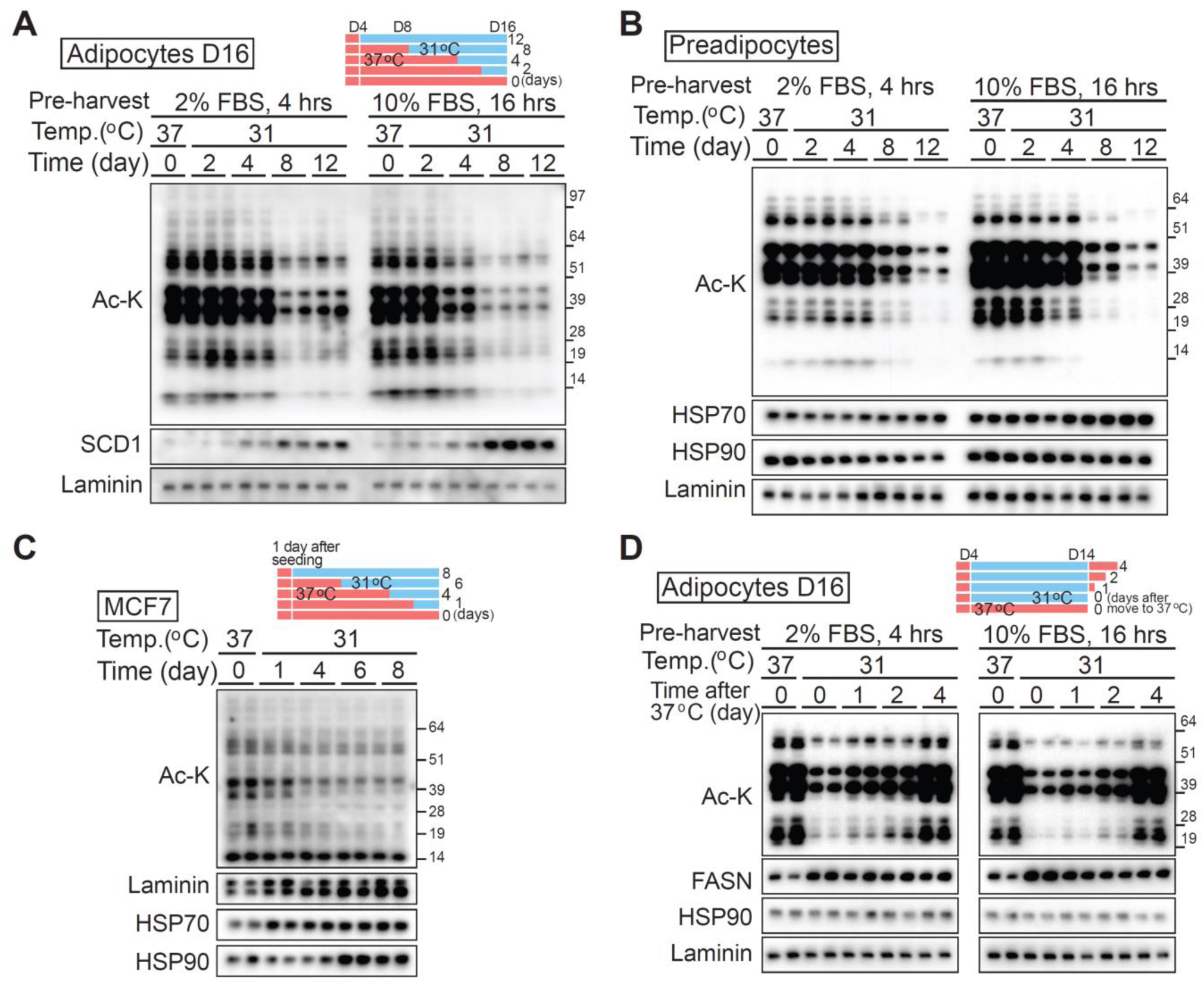
Cool adaptation decreases protein acetylation in precursors and differentiated adipocytes, independent of nutrient status. (**A** and **B**) Protein acetylation decreases in cool-adapted preadipocytes and differentiated adipocytes, regardless of nutrient status. (**A**) Cultured adipocytes and (**B**) preadipocytes were incubated at 31°C for the indicated durations. 2% FBS was used to minimize the influence of growth factors during the short-term incubation. Whole-cell lysates (WCL) were subjected to immunoblot analysis using antibodies against acetylated lysine. Laminin, HSP70, and HSP90 were used as loading controls. To assess the impact of nutrient status on protein acetylation, cells were cultured in medium containing either 2% FBS for 4 hours (left panel) or 10% FBS for 16 hours (right panel) prior to harvesting for both cell types. (**C**) Cultured adipocytes were incubated at either 37°C or 31°C for 10 days. Cells initially cultured at 31°C were returned to 37°C for the indicated durations to examine restoration of protein acetylation. WCL were subjected to immunoblot analysis using antibodies against acetylated lysine, FASN, HSP70, and laminin (loading controls). Nutrient conditions were as described in panels A and B. (**D**) The breast cancer cell line MCF7 was incubated at 31°C for the indicated durations. WCL were analyzed by immunoblotting for acetylated lysine, with laminin, HSP70, and HSP90 used as loading controls.

To determine whether the reduced protein acetylation is specific to adipocytes or represents a general cellular response, undifferentiated precursors were subjected to the same incubation conditions. These cells likewise exhibited substantially decreased protein acetylation at 31°C, regardless of the culture medium (**Figure 2B**). Thus, cool-induced hypoacetylation is a general cellular response and not confined to adipocytes.

To extend these findings, we also assessed breast-derived cell lines. Breast cancer cells, which exhibit increased metabolic activity and can generate localized heat [10; 11], were exposed to 31°C, a temperature range consistent with normal subcutaneous breast tissue conditions in healthy females [10]. This physiological window permits experimental exposure of breast cancer cell models to culture temperatures that mirror *in vivo* tissue environments outside the body core. As shown in Figure 2C and Supplemental Figure 1, exposure to 31°C resulted in a decrease in pan-acetylated protein levels, with timing and magnitude of hypoacetylation differing among cancer cell lines. Next, to determine if the decrease in protein acetylation was reversible, adipocytes were adapted to 31°C for 10 days, then switched back to 37°C. Acetylated protein levels returned to baseline within four days post-rewarming, independent of nutrient conditions, indicating that temperature-driven acetylation changes are both dynamic and reversible (**Figure 2D**).

Overall, exposure to cool temperatures reduces global protein acetylation in both mesenchymal stem cells and differentiated adipocytes, independent of nutrient status and reversible upon rewarming. Similar hypoacetylation can be induced in breast cancer cell models. Taken together, these findings highlight temperature as a dynamic and broadly conserved regulator of cellular protein acetylation.

### Mitochondria constitute the primary source of protein acetylation at 37 °C, which is globally reduced by cool adaptation

To characterize the subcellular localization of protein acetylation changes induced by cool adaptation, we differentiated adipocytes and incubated them at either 37°C or 31°C for 12 days (**Figure 3A**). Cells were fractionated into cytoplasmic and nuclear compartments using sequential hypotonic and hypertonic buffer extractions, and the amount of acetylated lysine was assessed by immunoblotting. Immunoblot analysis revealed marked decrease in acetylated protein levels within the cytoplasmic fraction of cool-adapted adipocytes compared to controls, while nuclear acetylation was further reduced (**Figure 3A**). These results suggest that cool adaptation primarily affects non-nuclear, cytoplasmic protein acetylation. To further determine the source of acetylated proteins, we isolated mitochondria from adipocytes cultured at 31°C or 37°C (**Figure 3B**). Comparison with whole-cell lysates (WCL) suggests that mitochondria are a major site of cool-induced hypoacetylation in adipocytes. Taken together, these experiments reveal that cool adaptation induces a selective decrease in cytosolic protein acetylation, predominantly originating from the mitochondrial compartment.

**Figure 3.**
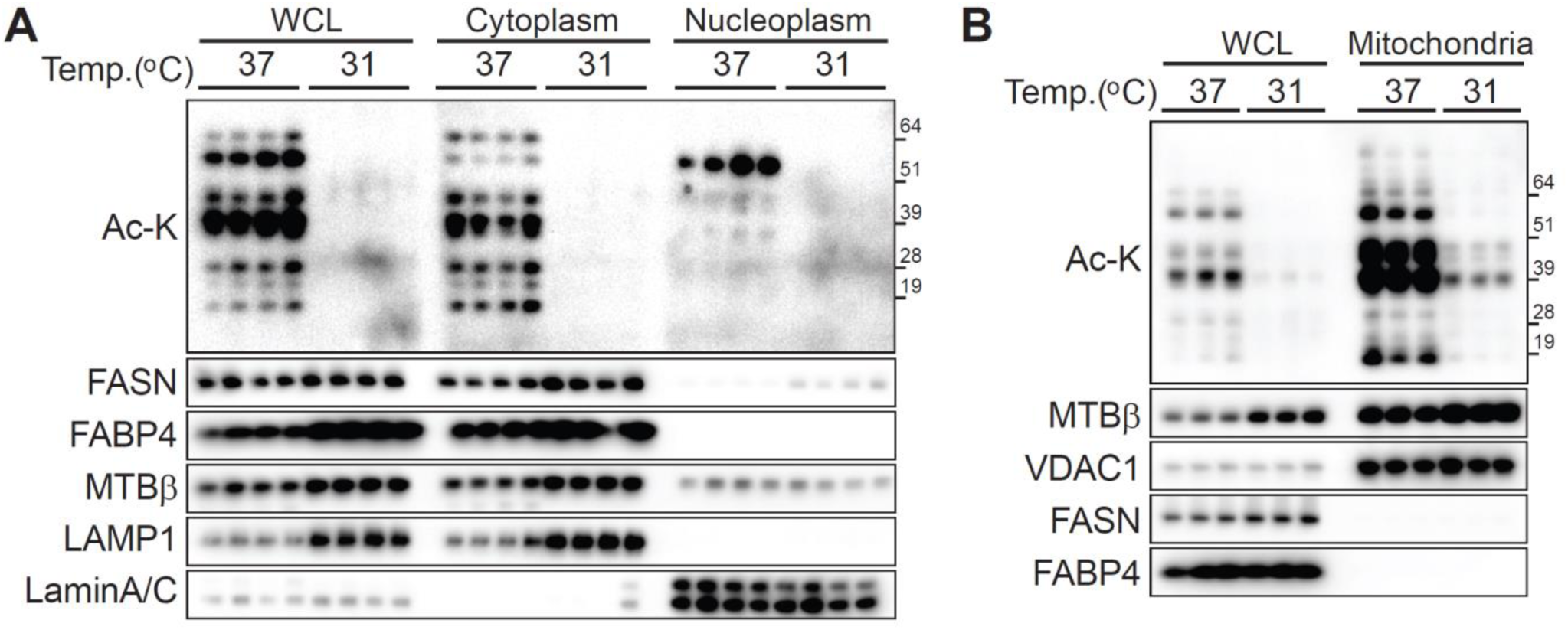
Cool adaptation decreases global protein acetylation, with mitochondria as the main source under control condition. (**A** and **B**) Cultured adipocytes were incubated at either 37°C or 31°C for 12 days. (**A**) After incubation, cells were separated into cytoplasmic and nuclear fractions using hypotonic and hypertonic buffers, respectively. Protein acetylation in these compartments and whole-cell lysates (WCL) was assessed by immunoblotting. Marker proteins included FASN (cytosol/ER), FABP4 (cytosol), MTBβ (mitochondria), LAMP1 (lysosome), and Lamin A/C (nuclear). Each sample was loaded at 14 µg. (**B**) Isolated mitochondria and WCL from adipocytes incubated under the same conditions as in panel A were analyzed by immunoblotting with the indicated antibodies to determine the subcellular distribution and sources of acetylated proteins. MTBβ and VDAC1 were used as mitochondrial markers. Each sample was loaded at 20 µg.

### Regulation of protein acetylation machinery in adipocytes under cool temperatures

To determine whether transcriptional changes in lysine acetyltransferases (KATs) or deacetylases (KDACs) - key regulators of protein acetylation - occur in response to cool adaptation, we quantified mRNA levels in adipocytes cultured at either 37°C or 31°C. Initial RNA-seq analysis indicated no significant changes in overall KAT or KDAC mRNA expression (NCBI GEO: GSE159451) [5]. However, targeted qPCR revealed that mRNA levels of several KATs (*Kat2b*, *Kat3a*, *Kat3b*, *Kat5*, *Kat6b*) were significantly decreased in adipocytes maintained at 31°C (**Figure 4A**). While this pattern aligns directionally with decreased global protein acetylation, these KATs are predominantly nuclear enzymes [12; 13], whereas our prior data demonstrated protein acetylation was mainly decreased in the mitochondrial fraction (**Figure 3**). Among KDACs, *Hdac11* and *Sirt3* exhibited slightly increased mRNA expression at 31°C (**Figure 4B**), potentially compatible with enhanced deacetylation [13].

**Figure 4.**
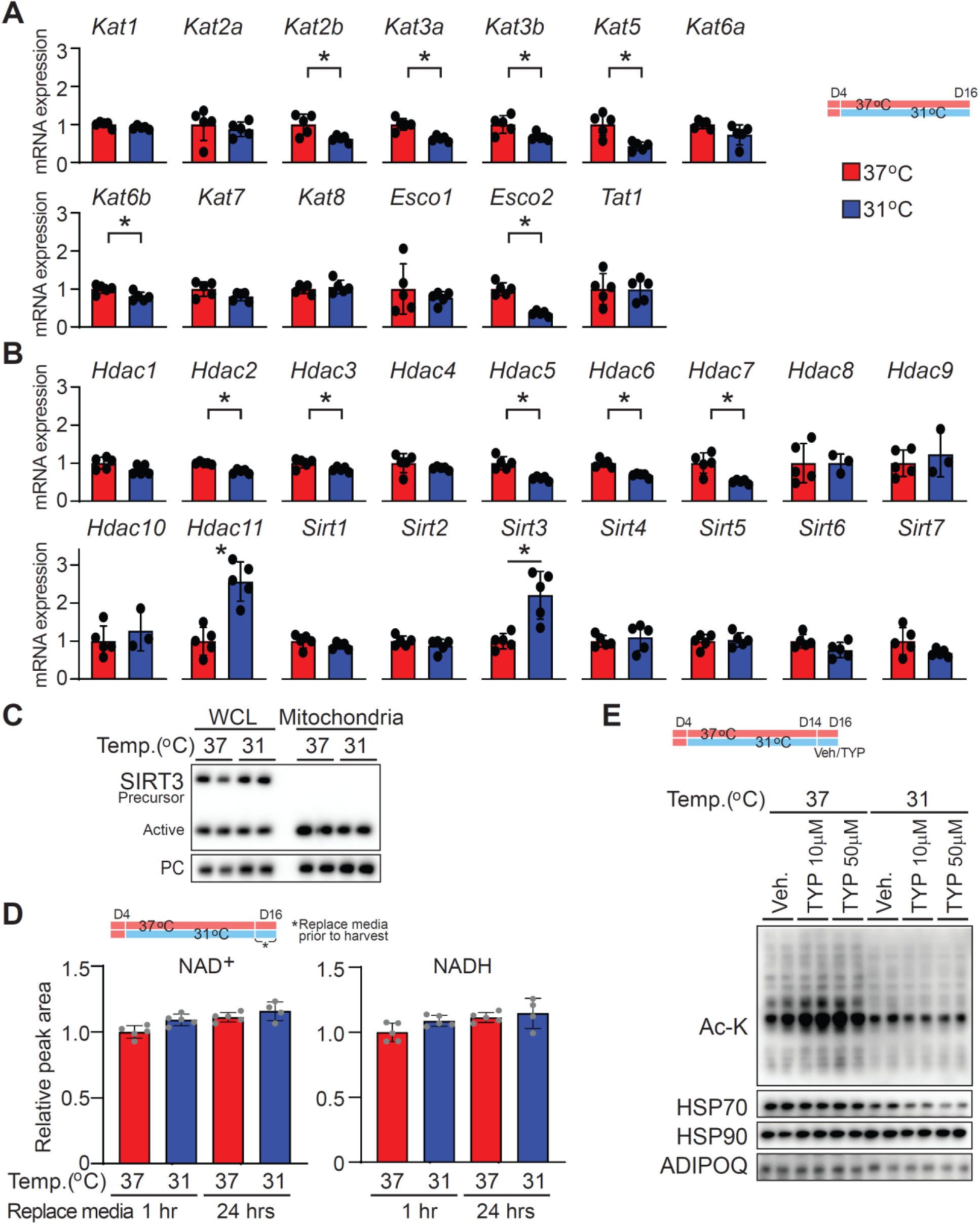
Regulation of protein acetylation machinery in adipocytes under cool temperatures. (**A** and **B**) Relative mRNA expression of lysine acetyltransferases (KATs) and deacetylases (KDACs). (**A**) Abundance of mRNAs coding for lysine acetyltransferases or (**B**) lysine deacetylases. Expression values were normalized to the geometric mean of the housekeeping genes *Hprt*, *Tbp*, *Gapdh*, and *Ppia*, and shown relative to the 37°C control group (*n* = 5 per group). (**C**) Isolated mitochondria and whole cell lysates (WCL) from adipocytes incubated at either 37°C or 31°C for 12 days were analyzed by immunoblotting. PC (Pyruvate carboxylase) served as the mitochondrial loading control, while both precursor (inactive) and mature (active) forms of Sirt3 were assessed as mitochondrial proteins of interest. Each sample was loaded at 20 µg. (**D**) Differentiated adipocytes were cultured at either 37°C or 31°C for 12 days and then incubated with fresh media for either one or 12 hrs prior to harvest. Metabolomic analyses of NAD⁺ and NADH concentrations was performed, and data are presented relative to the 37°C control at the 1-hour time point (*n* = 5 per group). Values represent mean ± SD. (**E**) Immunoblot analysis of pan–acetylated lysine in differentiated adipocytes cultured at either 37°C or 31°C for 12 days and then treated with the SIRT3 inhibitor 3-TYP (10 or 50 µM) for two days prior to harvest. HSP70, HSP90, and ADIPOQ were used as loading controls. Data are presented as mean ± SD. *p < 0.05.

Given SIRT3’s established role in mitochondrial protein deacetylation, we assessed expression of SIRT3 in both WCL and isolated mitochondria. The abundance of both precursor (inactive) and mature (active) forms of SIRT3 was similar between temperatures (**Figure 4C**), indicating that culture temperature does not affect SIRT3 protein levels. Additionally, NAD⁺ concentrations, the essential cofactor for SIRT3 activity, remained unchanged at both 1- and 24-hr time points, regardless of nutrient status (**Figure 4D**). To directly assess Sirt3’s involvement in regulating protein acetylation, adipocytes cultured at 37°C or 31°C for 12 days were treated with the SIRT3 inhibitor 3-TYP (10 or 50 µM) for 2 days (**Figure 4E**). While 3-TYP slightly increased pan-acetyl-lysine at 37°C, it failed to increase acetylation in 31°C cultures, suggesting SIRT3 inhibition cannot reverse cool-induced hypoacetylation. Taken together, these data demonstrate that cool adaptation reduces mitochondrial protein acetylation independently of KAT or KDAC expression, SIRT3 protein levels, NAD⁺ availability, or SIRT3 enzymatic activity, implicating alternative mechanisms such as altered acetyl-CoA flux or non-canonical regulatory processes.

### Modulation of acetyl-CoA metabolism does not alter protein acetylation in adipocytes at either 37°C or 31°C

Next, we investigated whether protein acetylation levels especially at 31°C could be increased in adipocytes by targeting specific metabolic pathways. Since acetyl-CoA is utilized during *de novo* lipogenesis [8; 14], we treated cells with the acetyl-CoA carboxylase (ACC) inhibitors TOFA and ND630 for two days, aiming to increase acetyl-CoA availability (**Figure 5A**). However, neither inhibitor affected protein acetylation at 37°C or 31°C. Acetate is converted to acetyl-CoA via acetyl-CoA synthetase short-chain family member (ACSS2) [15], so we next incubated cells with sodium acetate for two or four days. While protein acetylation levels increased at 37°C, no corresponding increase was observed at 31°C (**Figure 5B**). These results indicate that manipulating acetyl-CoA metabolism does not promote increased protein acetylation in adipocytes adapted to 31°C. To further explore the metabolic sources of acetylated proteins at 37°C, we considered mitochondrial acetyl-CoA production from glycolysis, fatty acid β-oxidation, and the catabolism of branched-chain amino acids - valine, leucine, and isoleucine [7; 16]. In glycolysis, mitochondrial pyruvate is converted to acetyl-CoA via the pyruvate dehydrogenase complex (PDC) [7; 16]. To probe the impact of pyruvate import, adipocytes were treated with the mitochondrial pyruvate carrier (MPC) inhibitor UK5099 at low and high concentrations for 2 days; however, neither treatment affected protein acetylation at 37°C or 31°C (**Figure 5C**). To evaluate the role of exogenous fatty acids in contributing to acetyl-CoA pools, cells were cultured in low concentrations of FBS and low glucose, supplemented with or without 100 μM non-esterified fatty acids (NEFAs; C16:0, C16:1, C18:1). Supplementation with NEFA had no effect on protein acetylation levels at 37°C or 31°C (**Figure 5D**). Furthermore, inhibition of long-chain fatty acid import into mitochondria using the carnitine palmitoyltransferase-1 (CPT1) inhibitor Etomoxir (10 or 20 μM) did not reduce protein acetylation at 37°C (**Figure 5E**). Additionally, supplementation with nicotinic acid (20-100 μM) to increase NAD+-dependent deacetylase activity did not affect protein acetylation levels at either temperature (**Supplemental Figure 2**). Taken together, these data demonstrate that manipulating acetyl-CoA synthesis or substrate availability from glucose, acetate, or fatty acids does not enhance protein acetylation in adipocytes adapted to 31°C. The robust acetylation observed at 37°C appears to depend on additional regulatory mechanisms beyond upstream substrate supply within the assessed metabolic pathways.

**Figure 5.**
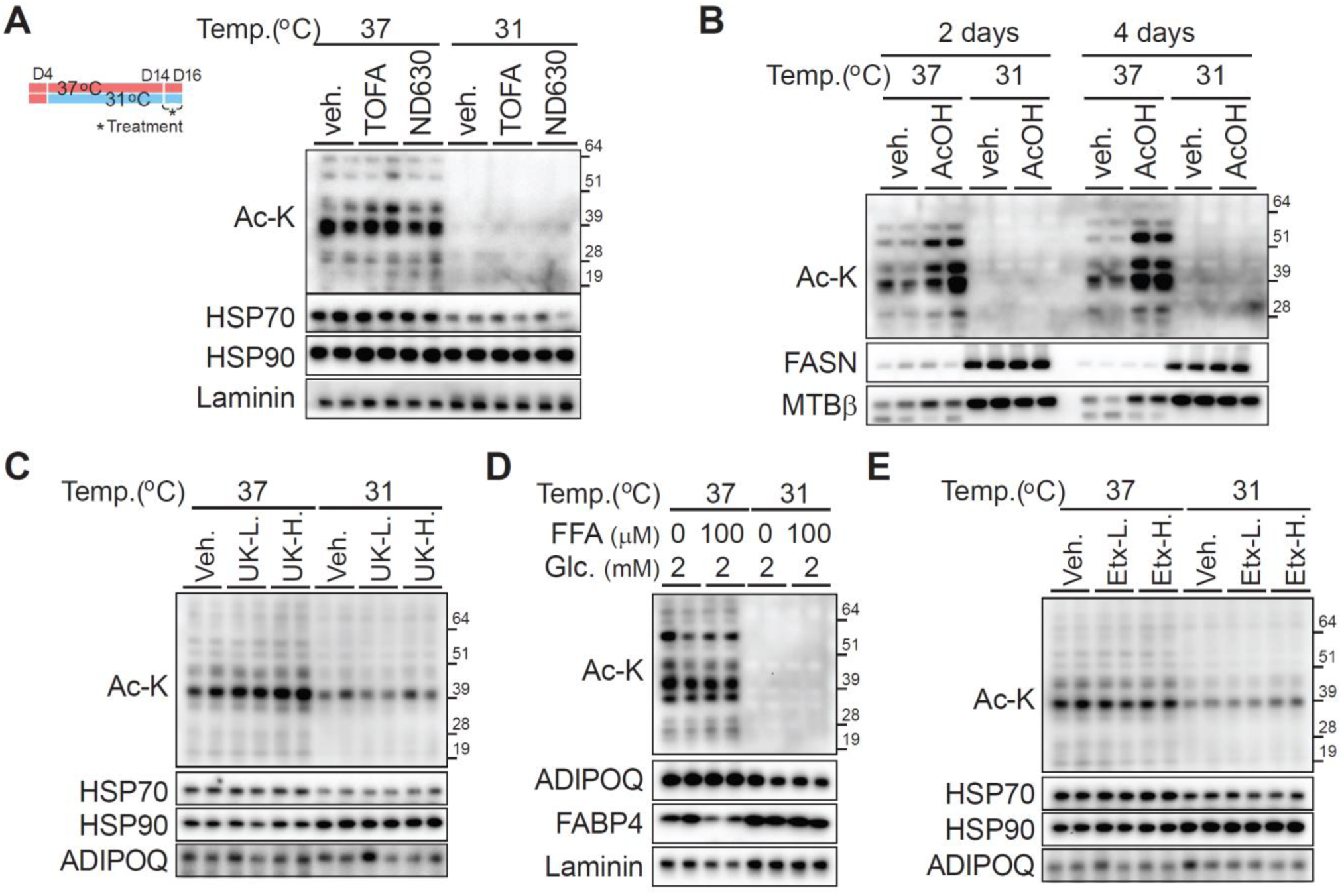
Exogenous sodium acetate increases protein acetylation at 37°C, but modulation of acetyl-CoA metabolism otherwise does not alter acetyl-lysine abundance in adipocytes at either 37°C or 31°C. (**A**–**E**) To modulate protein acetylation levels, differentiated adipocytes were cultured at either 37°C or 31°C for 10 days, and then treated with the indicated stimuli for two days, except in panel B, where cells were treated for either two or four days. Pan–acetylated lysine immunoblotting was used to assess acetylation, while separate blots served as loading controls. (**A**) Acetyl-CoA carboxylase inhibitors: TOFA (1 μM) and ND630 (0.2 μM). (**B**) Supplementation with 2 mM sodium acetate for 2 or 4 days. (**C**) Mitochondrial pyruvate carrier inhibitor UK5099 at either 2 or 10 μM. (**D**) 5% FBS and 2 mM glucose with or without 100 μM non-esterified free fatty acids (NEFAs; 33.3 μM each of C16:0, C16:1, and C18:1). (**E**) Etomoxir at 10 or 20 μM.

### Integrated proteomic and metabolomic profiling reveals temperature-dependent protein acetylation and metabolic changes in adipocytes

To identify which proteins undergo acetylation under the studied conditions and whether these modifications affect proteomic profiles, differentiated adipocytes were cultured at either 37°C or 31°C for 12 days. Immunoprecipitation (IP) of acetyl-lysine-modified peptides following trypsin digestion was performed (**Supplemental Data 1**), alongside metabolomic analysis (**Supplemental Data 2**). This combined approach enabled sensitive and large-scale identification and quantification of protein acetylation sites. Furthermore, we examined metabolite levels associated with enzymatic reactions regulated by acetylated proteins (**Supplemental Data 1**).

Specifically, among proteins highly acetylated at 37°C, we identified serine hydroxymethyltransferase 2 (SHMT2) and propionyl-CoA carboxylase alpha chain (PCCA) as examples of where substrate and product metabolite levels changed in coordination with the acetylation status of their regulating enzymes. Due to budget constraints, only one sample per temperature was analyzed; key acetylated proteins predicted by the proteomics data were independently validated by IP and immunoblotting (**Figure 6A**). Consistent with the IP acetyl-lysine-modified peptides data, acetylation of SHMT2 and PCCA was higher at 37°C (**Figure 6A**), whereas pyruvate carboxylase - also predicted to be highly acetylated at 37°C - showed no change in acetylation levels. Higher serine and lower glycine concentrations were detected at 37°C, suggesting that acetylation of SHMT2 may be important for enzymatic activity (**Figure 6B**). For PCCA, increased acetylation was associated with lower intracellular propionyl-carnitine concentrations (C3) and no change in acetyl-carnitine (C2) (**Figure 6C**), with the altered C3/C2 ratio supporting acetylation-dependent modulation of PCCA activity. These integrative proteomic and metabolomic data reveal a dynamic acetylome landscape influenced by temperature and highlight specific metabolic enzymes whose activities may be modulated by acetylation, providing new insights into temperature-dependent metabolic regulation in adipocytes.

**Figure 6.**
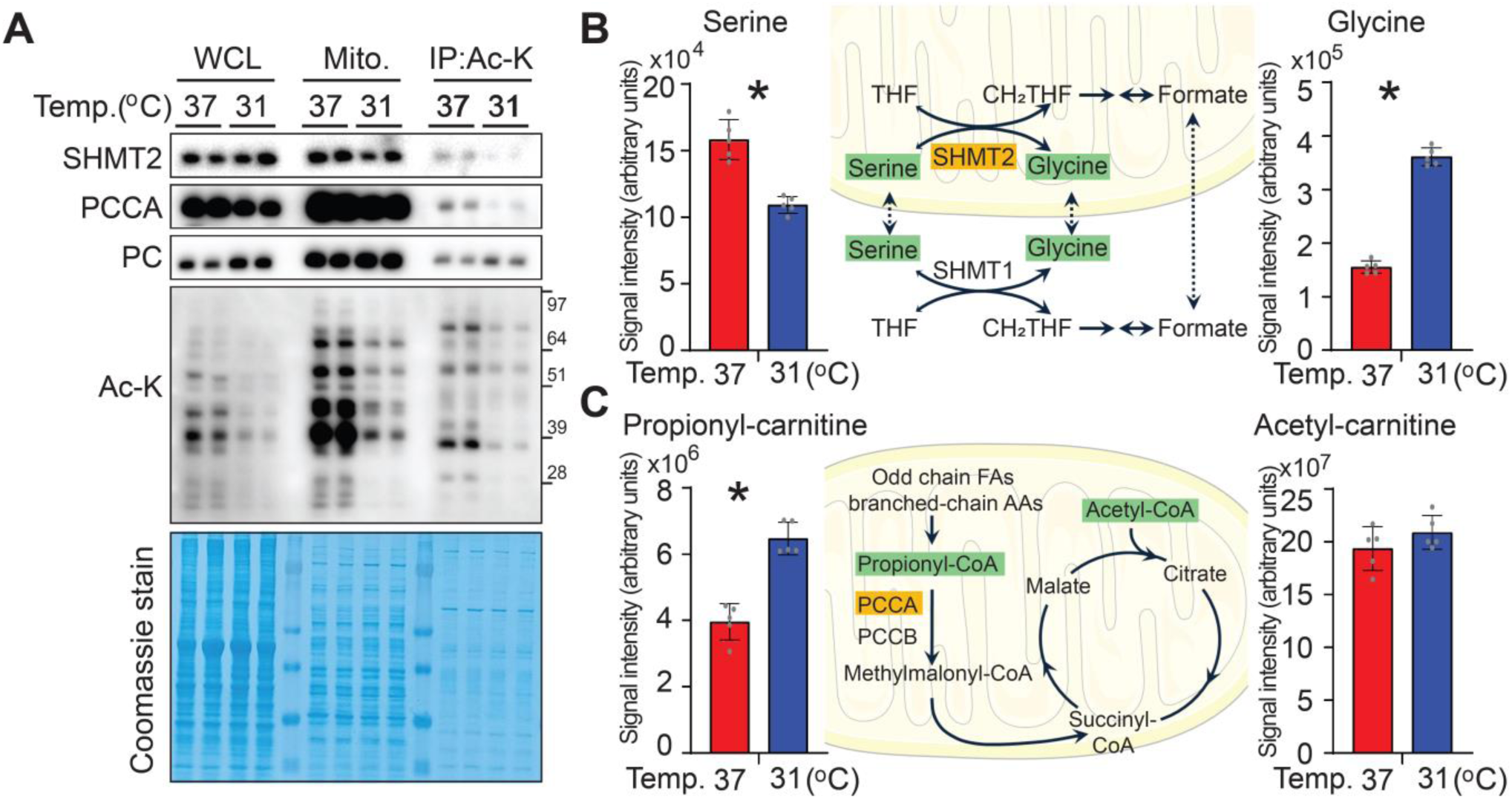
Integrated proteomic and metabolomic profiling reveals temperature-dependent protein acetylation and metabolic changes in adipocytes. (A) Validation of predicted acetylated proteins identified by acetyl-lysine peptide proteomics. Cultured adipocytes were incubated at either 37°C or 31°C for 12 days. Whole-cell lysates were subjected to immunoprecipitation (IP) with anti–acetyl-lysine antibodies, followed by immunoblotting for serine hydroxymethyltransferase 2 (SHMT2), propionyl-CoA carboxylase α (PCCA), and pyruvate carboxylase (PC), as predicted by proteomic analysis. (**B**-**C**) Metabolomic analyses corresponding to the acetylation status of target enzymes. Differentiated adipocytes were cultured at either 37°C or 31°C for 12 days and incubated with fresh media for 1 hour prior to harvest. Metabolite levels are presented as mean ± SD (*n* = 5 per group). (**B**) Increased serine and decreased glycine levels at 37°C were observed, correlating with elevated SHMT2 acetylation. (**C**) Elevated PCCA acetylation at 37°C was associated with decreased propionyl-carnitine (C3) and unchanged acetyl-carnitine (C2) levels. Data are presented as mean ± SD. *p < 0.05.

## DISCUSSION

### Potential mechanisms underlying temperature-dependent protein hypoacetylation

Our data reveal that global protein acetylation in adipocytes is significantly reduced during adaptation to cool temperatures (31°C), yet neither KAT/KDAC gene expression nor cofactor (acetyl-CoA) availability fully explain the effect. While RNA-seq revealed no significant changes in the overall expression, targeted qPCR detected decreased expression of several nuclear KATs (*Kat2b*, *Kat3a*, *Kat3b*, *Kat5*, *Kat6b*) at 31°C. Because these KATs are predominantly localized to the nucleus [12; 13], while the most pronounced loss of acetylation occurred in mitochondrial fractions, this spatial disparity indicates that transcriptional regulation of nuclear KATs is unlikely to be the major driver of mitochondrial hypoacetylation.

Among KDACs, *Hdac11* and *Sirt3* expression was slightly increased in cool-adapted cells, raising the possibility of enhanced deacetylation. However, total and mature SIRT3 protein were unaltered by temperature, and NAD⁺ concentrations - essential for sirtuin activity - remained constant. Notably, pharmacological inhibition of SIRT3 with 3-TYP elevated acetylation at 37°C but failed to do so at 31°C, indicating that SIRT3 activity alone is insufficient to reverse cool-induced hypoacetylation and that additional factors must be involved. This contrasts with Sirt3 knockout models, where SIRT3 loss triggers widespread mitochondrial hyperacetylation [17; 18]. The absence of a similar “deacetylation signature” in cool-adapted adipocytes, despite unchanged SIRT3 protein levels, suggests that temperature engages different mechanisms.

These findings are consistent with previous reports suggesting that protein acetylation is sensitive to acetyl-CoA compartmentalization and flux, not just KAT/KDAC transcription [6; 19; 20]. In hepatocytes and neurons, changes in acetyl-CoA levels can modulate mitochondrial and global protein acetylation independent of acetyltransferase or deacetylase abundance. In our model, cool adaptation likely alters acetyl-CoA flux or transport, constraining substrate availability for mitochondrial acetylation. Competing lysine modifications (e.g., lipoylation) may further influence acetylation under different thermal/metabolic states.

Despite extensive profiling, the precise mechanisms linking lower temperature to reduced protein acetylation remain unclear. Our data suggest non-canonical regulation, such as post-translational modification of acetylation machinery, altered subcellular localization, or acetyl-CoA metabolism, may underlie the observed hypoacetylation. Temperature-sensitive protein interactions and feedback from metabolic pathways may also affect acetylation, independent of mRNA or protein abundance.

### Temperature regulation and protein acetylation

SIRT3 is highly expressed in brown adipose tissue (BAT) and is upregulated by cold exposure [20]. Sebaa et al. demonstrated that Sirt3 knockout (KO) mice show increased mitochondrial protein acetylation in BAT - including hyperacetylation of UCP1 - and experience heat intolerance due to impaired thermogenesis [17]. Thus, SIRT3 appears to regulate thermogenic capacity mainly through deacetylation of upstream metabolic pathways, rather than direct control of UCP1, since mutation of SIRT3-targeted acetylation sites on UCP1 did not alter leak respiration.

These findings align with our observation that environmental temperature modulates global protein acetylation. In our study, exposing differentiated adipocytes to cooler temperatures resulted in a marked decrease in total protein acetylation. In contrast, BAT mitochondria from *Sirt3* KO and cold-exposed mice showed increased acetylation [17]. The key distinction lies in context: in our *in vitro* system, cool adaptation produced a generalized reduction in acetylation in both MSCs and differentiated adipocytes, independent of nutrient status, while *in vivo* cold exposure increased mitochondrial acetylation in BAT from approximately 1.4-fold at 22 °C to 1.8-fold at 4 °C [17], likely reflecting the locally warmer, highly respiring state of thermogenic tissue. In *Sirt3* KO skeletal muscle, mitochondrial protein acetylation is increased, but only for a subset of bands and to a lesser extent than in brown adipose tissue [17; 18]. One possible explanation is that interscapular BAT is locally warmer than skeletal muscle even at 22°C housing, due to ongoing UCP1-dependent thermogenesis, whereas skeletal muscle behaves as a cooler ‘shell’ tissue. If acetylation scales with local temperature and acetyl-CoA flux, this disparity between BAT and muscle in the same animals further supports the concept that tissue temperature is a critical determinant of mitochondrial protein acetylation.

Overall, these observations support the idea that mitochondrial protein acetylation is dynamically regulated by temperature and metabolic flux, and that the “functional” tissue temperature may differ from ambient temperature. Our work emphasizes broad changes in acetylation driven by extracellular temperature, largely independent of SIRT3 abundance or activity in cultured cells, whereas the BAT study highlights that mitochondrial deacetylation is essential for maintaining activity of electron transport chain and fatty acid oxidation proteins, as well as thermogenic capacity.

### Comparison with cancer studies on SHMT2 and PCCA acetylation

Among proteins highly acetylated at 37°C, we identified serine hydroxymethyltransferase 2 (SHMT2) and propionyl-CoA carboxylase alpha chain (PCCA) as examples where substrate and product metabolite levels changed in coordination with the acetylation status of their regulating enzymes (**Figure 6**). A recent study reported that SIRT3 promotes colorectal cancer by targeting SHMT2 [21]. In colorectal cancer cells, acetylation of SHMT2 at K95 inhibits its enzymatic activity, reducing serine-to-glycine conversion and one-carbon metabolism [21]. Maintaining SHMT2 K95 acetylation slows proliferation, while SIRT3-mediated deacetylation restores SHMT2 activity and promotes tumor growth [21]. Consistent with this, SHMT2 K95 acetylation is significantly lower in human colorectal carcinoma samples than in adjacent normal tissue [21]. A complementary study in gastric cancer cells showed functional SHMT2 acetylation at K200 [22]. Quantitative acetyl-proteomics in MKN-45 cells treated with an anticancer bioactive peptide–selenium chelate (ACBP-Se) revealed significant remodeling of the acetylome, including regulated acetylation of SHMT2 at K200 [22]. PRM validation confirmed changes in SHMT2 K200 acetylation, and pathway analysis linked this modification, along with others, to amino acid metabolism, one-carbon pathways, and redox processes associated with the antiproliferative effects of ACBP-Se [22]. Our results differ from these cancer studies mainly in the specific acetylation sites identified. Proteomics analysis after IP revealed that K147, K13, K42, K320, and K330 as highly acetylated in adipocytes at 37°C compared to 31°C. Although these sites differ from K95 [21] and K200 [22], a shared theme persists: SHMT2 acetylation is associated with reduced enzymatic activity and altered one-carbon metabolism, suggesting site-specific and context-dependent regulation. Sequence alignment shows that human and mouse SHMT2 share 92% identity, with nearly all lysine residues conserved. Human K95 and K200 - key acetylation sites in colorectal and gastric cancer - correspond to analogous lysines in mouse SHMT2, while the temperature-sensitive sites we detected (K13, K42, K147, K320, K330) are also located in highly conserved regions. This evolutionary conservation supports a broadly shared SHMT2 acetylation mechanism, though the specific lysines modified may vary by species, cell type, or physiological state.

Propionyl-CoA carboxylase (PCC), composed of the alpha (PCCA) and beta (PCCB) subunits, is the key mitochondrial enzyme that converts propionyl-CoA to methylmalonyl-CoA, thereby linking odd-chain fatty acid and branched-chain amino acid catabolism to the TCA cycle [23–25]. While PCC disease mutations are well characterized [23–25], physiological roles of PCCA lysine acetylation remain largely unexplored. Large-scale acetyl-proteomics datasets list PCCA as an acetylated mitochondrial protein; however, to our knowledge, no prior study has systematically mapped PCCA acetylation sites or tested how acetylation affects PCC activity or propionate flux. Our data, though based on one sample per condition, suggest PCCA is more highly acetylated at 37°C on lysines K61 and K184, coincident with decreased propionyl-carnitine (C3) compared to acetyl-carnitine (C2). This raises the possibility that PCCA acetylation is temperature-sensitive and influences propionyl-CoA utilization. Taking together with SHMT2 results, these observations support a broader model in which mitochondrial enzyme acetylation may fine-tune metabolic flux in response to environmental temperature, though further work is needed to clarify mechanisms, especially for PCCA.

## Method Details

### Animals

Three-week-old C57BL/6J male mice (The Jackson Laboratory, Ellsworth, ME, USA) were housed at either 22°C or at 30°C for 12 weeks. Mice were single housed with minimal bedding and no nesting materials. All animal studies were performed in compliance with policies of the University of Michigan Institutional Animal Care and Use Committee. The protocol number is PRO00009687.

### Adipocyte fractionation for bioenergetic assays

The psWAT depots were excised, minced with blades, and digested with 533.3 U/mL collagenase type I (LS004197, Worthington). Digestion was done in Hank’s Balanced Salt Solution (HBSS) (14025-092, Gibco) supplemented with 1 g/L glucose and 500 nm adenosine for 30 minutes at 37°C shaking horizontally at 100 rpm. Adipocytes were separated from the stromal vascular fraction by filtering through 300 μm filters and centrifuging at 100 x g for 8 minutes. Buoyant adipocyte fractions were transferred to 5 ml tubes and washed 3 times with STE-BSA buffer (250 mM sucrose, 5 mM Tris, 2 mM EGTA, 4% fatty acid free BSA, pH 7.4) by removing the infranatant from beneath the adipocyte layer with a 5 mL syringe and 21G long needle. Bioenergetic parameters of floated adipocytes were then assessed with an Oroboros-2k.

### Respirometry assay of adipocytes

Oxygen consumption of floated adipocytes was measured with high-resolution respirometer using an Oxygraph-2k (OROBOROS Instruments, Innsbruck, Austria). Floated adipocytes (40 μL), freshly isolated from psWAT, were pipetted into 2 mL mitochondrial respiration medium buffer (MIR05) with the stirrer set to 750 rpm. The bioenergetic parameters measured were basal respiration (5 mM pyruvate, 1 mM malate, 5 mM succinate), proton leak respiration (1 μM oligomycin), and maximal respiration (1 μM FCCP steps until we observed a plateau in oxygen consumption). Non-mitochondrial oxygen consumption (2.5 μM antimycin A) was subtracted from the other respiratory states. ATP-linked respiration was calculated by subtracting leak respiration from baseline, and spare capacity was calculated by subtracting baseline from maximal respiration. Oxygen consumption rates were normalized to total genomic DNA isolated from adipocyte suspensions. Samples were homogenized in lysis buffer (50 mM Tris-HCL, 0.45% NP-40, 0.45% Tween-20, 0.2 mg/mL Proteinase K) and incubated on a heat block at 55°C for 2 hours shaking at 700 rpm. After proteinase K inactivation, RNase A (0.5 mg/mL) was added, and samples were incubated at 37°C for 10 minutes. For DNA precipitation, absolute ethanol was added and samples were centrifuged at 12000 x g for ten minutes. The resultant pellets were washed twice with 70% ethanol, air-dried, and resuspended with nuclease free water. Total genomic DNA was quantified using a NanoDrop spectrophotometer.

### Isolation and differentiation of adipocyte precursors

Adipocyte precursors were isolated from ears of mice of the indicated genotypes, as previously described [5; 26]. Cells were maintained in 5% CO_2_ and DMEM/F12 1:1 media (Gibco; Invitrogen 11330-032) supplemented with 10% FBS (Cytiva HyClone), primocin (InVivoGen), and 10 ng/ml recombinant bFGF (PeproTech). For induction of adipogenesis, recombinant bFGF was removed and replaced with 10% FBS containing 0.5 mM methylisobutylxanthine (Cayman Chemical 13347), 1 μM dexamethasone (Sigma-Aldrich D1756), 5 μg/ml insulin (Sigma-Aldrich I3536), and 5 μM rosiglitazone (Cayman Chemical 71740). On day 2, cells were fed 5 μg/ml insulin plus 5 μM troglitazone. On day 4 and every 2 days thereafter, cells were fed with 10% FBS-supplemented media. All the media conditions used in the experiments within this study contain 10% FBS, unless otherwise specified in the individual figure legends.

### Nuclear and Cytoplasmic Fractionation

Nuclear and cytoplasmic fractions were isolated using the NE-PER™ Nuclear and Cytoplasmic Extraction Reagents (Thermo Fisher Scientific) according to the manufacturer’s protocol. Briefly, cells were harvested and washed with ice-cold PBS, then resuspended in Cytoplasmic Extraction Reagent I (CER I) supplemented with protease/phosphatase inhibitors plus deacetylase inhibitors (50 µM EX-527, 10 µM trichostatin A; TSA, 10 mM sodium butyrate, 5 µM Vorinostat; SAHA) and incubated on ice. Cytoplasmic Extraction Reagent II (CER II) was added, samples were vortexed briefly, and cytoplasmic lysates were collected after high-speed centrifugation to pellet intact nuclei. The remaining nuclear pellets were resuspended in Nuclear Extraction Reagent (NER) containing protease/phosphatase/deacetylase inhibitors, incubated on ice with intermittent vortexing, and nuclear extracts were collected after centrifugation

### Isolation of mitochondria from cultured MSC adipocytes

Isolation of mitochondria was as described previously [5]. Briefly, adipocytes from culture or isolated from gluteal WAT were homogenized using a Potter-Elvehjem homogenizer and centrifuged at 800 x g for 10 minutes at 4°C. The supernatant was then centrifuged for 15 minutes at 8,000 x g at 4°C, and the pellet was washed with ice-cold buffer. After centrifugation at 7,000 x g for 10 minutes at 4°C, the pellet containing mitochondria was resuspended for analyses.

### LC-MS metabolomics

For metabolomics assays, adipocytes were cultured in 6-well plates and treated as described in the manuscript. When cells were ready for extraction, all wells were aspirated of media, rapidly rinsed with 150 mM ammonium acetate to remove residual extracellular salts and metabolites, rapidly aspirated again, and frozen by direct addition of liquid nitrogen to the wells. The entire rinsing process was accomplished within ∼20 s to rapidly quench metabolism and minimize metabolic perturbations. The plates were maintained at -80°C until the day of extraction, at which point 300 µL of ice-cold 9:1 methanol: chloroform was added to each well. Each well was scraped individually with a cell scraper in the presence of the solvent for approximately 10 seconds per well to release and lyse cells. Cell debris and solvent were transferred to labeled polypropylene microcentrifuge tubes on wet ice for an additional 10 min and then centrifuged at 20,000 x g at 4°C for 10 minutes. 180 µL of supernatant from each well was transferred to LC autosampler vials containing micro-inserts and the solvent was evaporated to dryness using a gentle stream of room-temperature nitrogen gas. The samples were reconstituted in 50 µL of 80:20 water: methanol, vortexed, and submitted to LC-MS analysis. A QC sample was generated by pooling residual extract from the samples and drying and reconstituting at the same ratio as the samples; this sample was analyzed as every 10th run in the sample queue.

Ion paring liquid chromatography-mass spectrometry was performed on an Agilent system consisting of a 1290 UPLC module coupled with 6530 Quadrupole-Time-of-flight (QTOF) mass spectrometer (Agilent Technologies, Santa Clara, CA.) using a JetStream ESI source in negative mode. The UPLC was equipped with a 10-port valve configured to allow the column to be either eluted to the mass spectrometer or back-flushed to waste. The chromatographic separation was performed on an Agilent ZORBAX RRHD Extend 80Å C18, 2.1 × 150 mm, 1.8 μm column with an Agilent ZORBAX SB-C8, 2.1 mm × 30 mm, 3.5 μm guard column. Mobile phase A consisted of 97:3 water/ methanol and mobile phase B was 100% methanol; both A and B contained tributylamine and glacial acetic acid at concentrations of 10 mM and 15 mM, respectively. The column was back-flushed with mobile phase C (100% acetonitrile, no additives) between injections for column cleaning. The LC gradient was as follows: 0-2 min, 0%B; 2-11 min, linear ramp from 0-99%B; 12-16 min, 99%B, 16-17.5min, 99-0%B. At 17.55 min, the 10-port valve was switched to reverse flow (back-flush) through the column. From 17.55-20.45 min the solvent was ramped from 99%B to 99%C. From 20.45-20.95 min the flow rate was ramped up to 0.8 mL/min, which was held until 22.45 min, then ramped down to 0.6mL/min by 22.65 min. From 22.65-23.45 min the solvent was ramped from 99% to 0% C while flow was simultaneously ramped down from 0.6-0.4mL/min. From 23.45 to 29.35 min the flow was ramped from 0.4 to 0.25 mL/min; the 10-port valve was returned to restore forward flow through the column at 25 min. An isocratic pump was used to introduce reference mass solution through the reference nebulizer for dynamic mass correction. Total run time was 30 min. Compounds were identified by matching accurate mass and retention time with authentic standards analyzed on the same instrument. Relative quantitation by peak area was performed using Agilent Quantitative Analysis software.

Reversed phase LC-MS analysis of samples were performed using a Thermo Vanquish Horizon UPLC system coupled to an Orbitrap ID-X mass spectrometer operated in positive ion mode. The column used was a Waters HSS T3 100 mm x 2.1mm inner diameter with 1.8 µm particle size with a matched Vanguard guard column. Mobile phase A was 0.1% formic acid in water and mobile phase B was 0.025% formic acid in methanol. The gradient consisted of a linear ramp from 0 to 100% B over 10 minutes, a 6-minute hold at 100% B, after which the mobile phase returned immediately to 0% B and was held for 3 minutes to re-equilibrate the column prior to the next run. The column temperature was set at 55 °C, the flow rate was 0.45 mL/min, and the injection volume was 10 µL. MS source parameters were as follows. Spray voltage: 3500V positive / 2500V negative, sheath gas: 50, aux gas: 10, sweep gas: 1, ion transfer tube temperature: 325 °C, vaporizer temperature: 350 °C. MS scan parameters were as follows. Scan mode: MS1 full-scan, orbitrap resolution: 120,000, quadrupole isolation: enabled, scan range: m/z 100-1000, RF lens: 50%, ACG target: standard, Maximum injection time mode: Auto, data type: centroid, Easy-IC (internal calibration): enabled. Feature detection and relative quantitation was performed with MSDIAL v5 software [27]. Identities of compounds reported in the manuscript were confirmed by analysis of authentic standards.

### Acetyl-proteome

Acetyl-proteome analysis was performed by PTM Bio (Chicago, IL); the method is briefly described as follows: Dried S-Trap™ elutions were reconstituted with 1 mL Desalting Buffer (1% ACN, 0.1% TFA, pH < 2), samples’ pH was adjusted further with TFA until pH < 2 when necessary. Samples were then loaded, desalted, and eluted out of 50 mg/ml HyperSep™ C18 Cartridge. The eluted samples were frozen and dried via SpeedVac. Dissolving the peptide segments in IP buffer solution (100 mM NaCl, 1 mM EDTA, 50 mM Tris-HCl, 0.5% NP-40, pH 8.0), transfer the supernatant to a lactated resin that has been pre-washed (PTM104), sourced from Hangzhou Jingjie Biotechnology Co., Ltd., PTM Bio), and place it on a rotary shaker in a 4°C environment, gently shaking and incubating overnight. After the incubation, wash the resin four times with IP buffer solution and twice with deionized water. Finally, use 0.1% trifluoroacetic acid eluent to elute the peptide segments bound to the resin. Elute three times in total, collect the eluate and freeze-dry it in a vacuum. After freeze-drying, remove salt according to the instructions of C18 ZipTips, and then provide it for liquid chromatography-mass spectrometry analysis.

### Detection of specific proteins by immunoblot

After lysis in 1% NP-40, 120 mM NaCl, 50 mM Tris-HCl (pH 7.4), 50 mM NaF, 2 mM EDTA, 1× protease inhibitor cocktail (Sigma-Aldrich), deacetylase inhibitors (50 µM EX-527, 10 µM TSA, 10 mM sodium butyrate, 5 µM SAHA), protein concentrations of lysates after centrifugation were measured by BCA protein assay (Thermo Fisher Scientific). Lysates were diluted to equal protein concentrations in lysis buffer and then boiled in SDS sample buffer (20 mM Tris; pH 6.8, 2% SDS, 0.01% bromophenol blue, 10% glycerol, 5% 2-mercaptoethanol) and subjected to SDS-PAGE and immunoblotting according to standard techniques [28].

### Immunoprecipitation

Mitochondrial lysates were pre-cleared with Protein A/G magnetic beads (Cell Signaling Technology #73778) to reduce nonspecific binding. Briefly, beads were resuspended by gentle vortexing, then 20 μl of bead slurry was transferred to a clean tube, placed on a magnetic rack, and the storage buffer was removed. Beads were washed twice with 500 μl of 1× cell lysis buffer, and 20 μl of pre-washed beads was added to 200 μl of mitochondrial lysate. The lysate was incubated with rotation for 20 min at room temperature, and the pre-cleared lysate was transferred to a clean tube.

For immunoprecipitation, anti-acetyl-lysine antibody (Cell Signaling Technology #9441) was added to 200 μl of pre-cleared lysate at the dilution recommended by the manufacturer and incubated overnight at 4°C with rotation to allow immunocomplex formation. Pre-washed Protein A/G magnetic beads were then added, and samples were incubated with rotation for 60 min at room temperature. Beads were collected on a magnetic rack and washed five times with 500 μl of 1× cell lysis buffer. Bound proteins were eluted in 3× SDS sample buffer, heated at 95°C for 5 min, and the supernatant was collected for SDS-PAGE and immunoblot analysis. To minimize detection of IgG heavy and light chains, immunoblotting was performed using Mouse Anti-Rabbit IgG (Conformation Specific) Antibody (HRP Conjugate; Cell Signaling Technology #5127) as the secondary antibody.

### mRNA quantification by RT-PCR

RNA isolation, reverse transcription, and quantitative PCR were performed as previously described [28].

### Statistics

All data are presented as mean ± SD. When comparing 2 groups, significance was determined using 2-tailed Student t test. When comparing multiple experimental groups, an analysis of variance (ANOVA) was followed by post hoc analyses with Dunnett or Sidak test, as appropriate. Differences were considered significant at p < 0.05 and are indicated with asterisks. For metabolomics data analyses, the proportion of labeling at each carbon position was calculated by dividing each species by total sum of peak areas of all labeled positions.

The proportion of data is well known to follow a beta distribution. Beta regression model is an extension of the generalized linear model with an assumption that the response variable follows a beta distribution with values in standard unit interval (0,1).

## Data Availability

Data files for proteomics and metabolomics containing tabulated feature metadata and quantitative values are included as supplemental data section of the manuscript this paper.

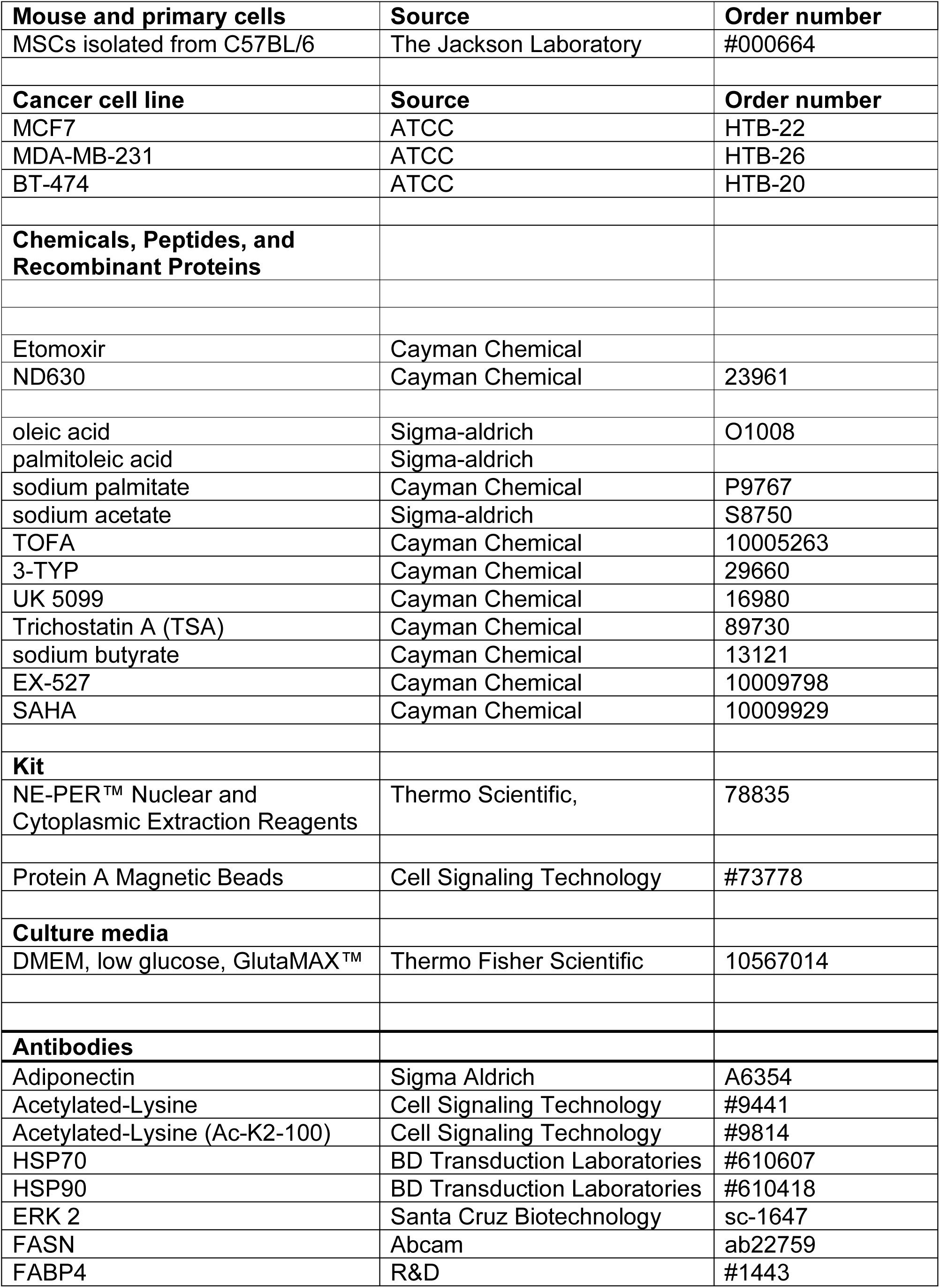
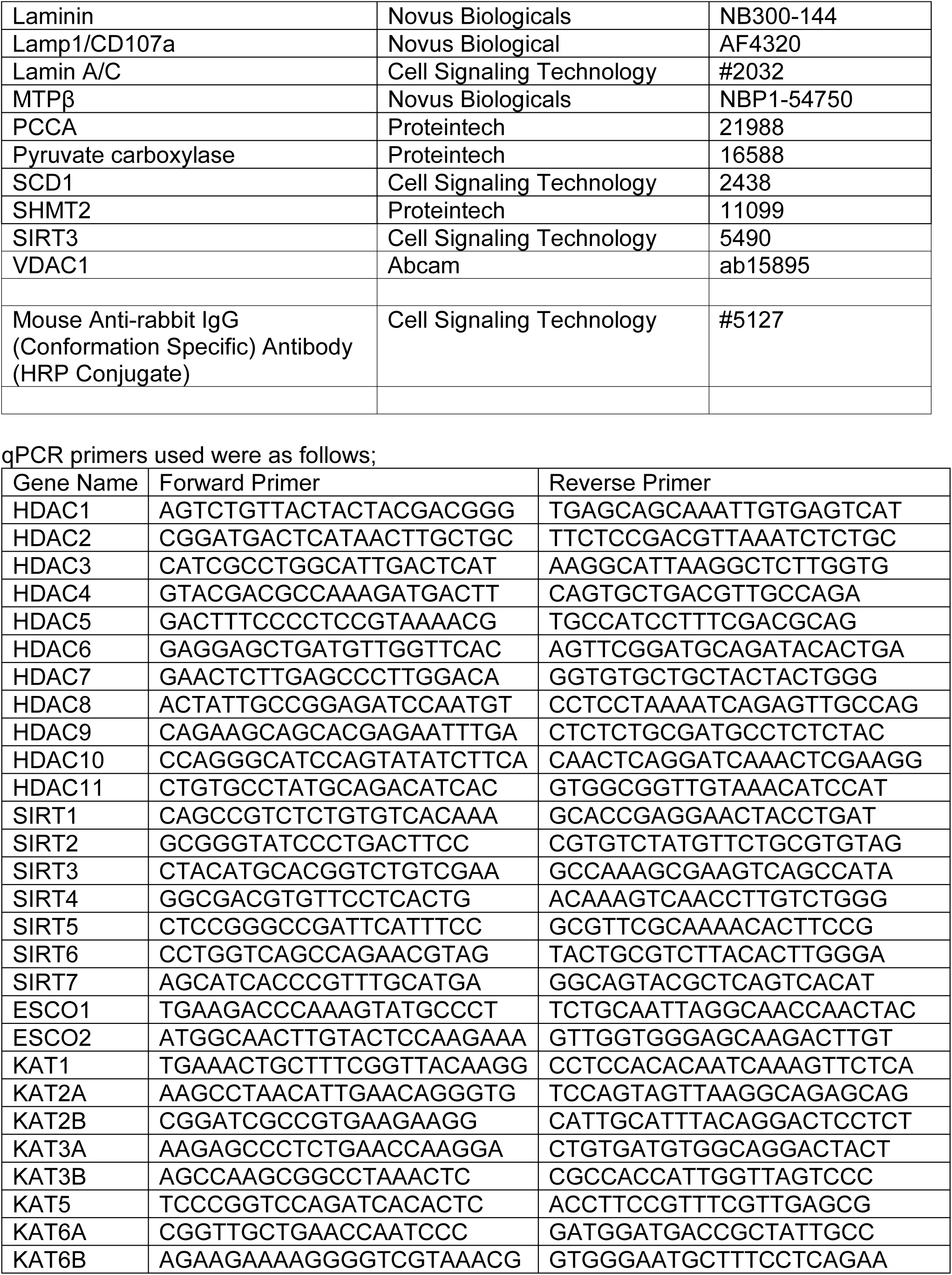
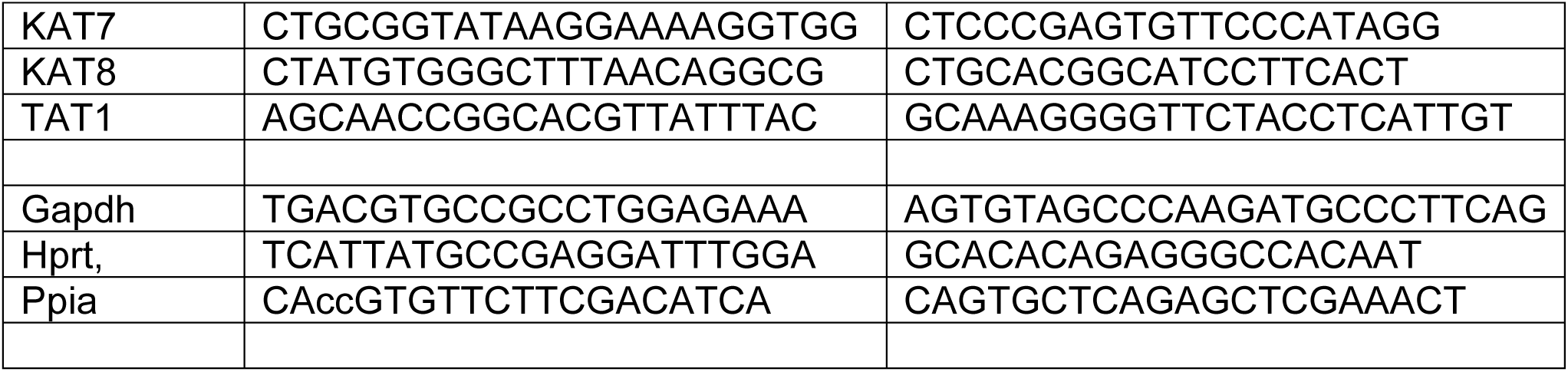

## ACKNOWLEDGEMENTS

This work was supported by grants or fellowships from the NIH to OAM (RO1 DK121759; R01 DK125513, and R01 DK130879). This research was also supported by core facilities of the Michigan Diabetes Research Center (P30 DK020572), Michigan Nutrition and Obesity Center (P30 DK089503), and University of Michigan Frankel Cardiovascular Regeneration Core Laboratory. We gratefully acknowledge Dr. Dan Beard and Françoise Van den Bergh at the University of Michigan for providing access to the Oxygraph-2k instrument used for bioenergetic measurements. We also thank Lanna Lewis and Aida Machuca for technical assistance, Guanghui Han for discussions regarding proteomics data analysis, and members of the MacDougald lab for helpful discussions and assistance.

