## Supplemental Figures for "Adaptation of white adipocytes to cooler temperatures: impacts on energy metabolism and protein acetylation"

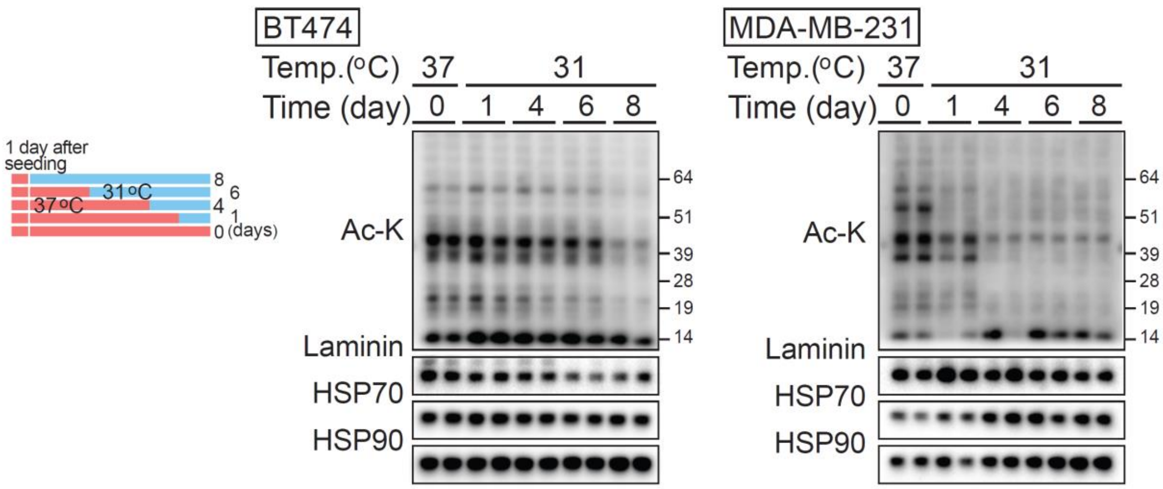


**Supplemental Figure 1.** The breast cancer cell line BT474 and MDA-MB-231 was incubated at 31°C for the indicated durations. Whole-cell lysates were analyzed by immunoblotting for acetylated lysine, with laminin, HSP70, and HSP90.

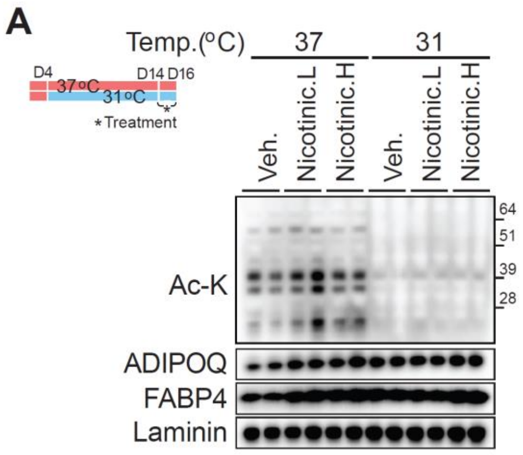


**Supplemental Figure 2.** Differentiated adipocytes were cultured at either 37°C or 31°C for 10 days, followed by treatment with 20–100 μM nicotinic acid for 2 days.
